# Co-transcriptional folding of a bio-orthogonal fluorescent scaffolded RNA origami

**DOI:** 10.1101/864678

**Authors:** Emanuela Torelli, Jerzy W. Kozyra, Ben Shirt-Ediss, Luca Piantanida, Kislon Voïtchovsky, Natalio Krasnogor

## Abstract

The scaffolded origami technique has provided an attractive tool for engineering nucleic acid nanostructures. This paper demonstrates scaffolded RNA origami folding *in vitro* in which all components are transcribed simultaneously in a single-pot reaction. Double-stranded DNA sequences are transcribed by T7 RNA polymerase into scaffold and staple strands able to correctly fold in high yield into the nanoribbon. Synthesis is successfully confirmed by atomic force microscopy and the unpurified transcription reaction mixture is analyzed by an in gel-imaging assay where the transcribed RNA nanoribbons are able to capture the specific dye through the reconstituted split Broccoli aptamer showing a clear green fluorescent band. Finally, we simulate the RNA origami *in silico* using the nucleotide-level coarse-grained model oxRNA to investigate the thermodynamic stability of the assembled nanostructure in isothermal conditions over a period of time.

Our work suggests that the scaffolded origami technique is a valid, and potentially more powerful, assembly alternative to the single-stranded origami technique for future *in vivo* applications.

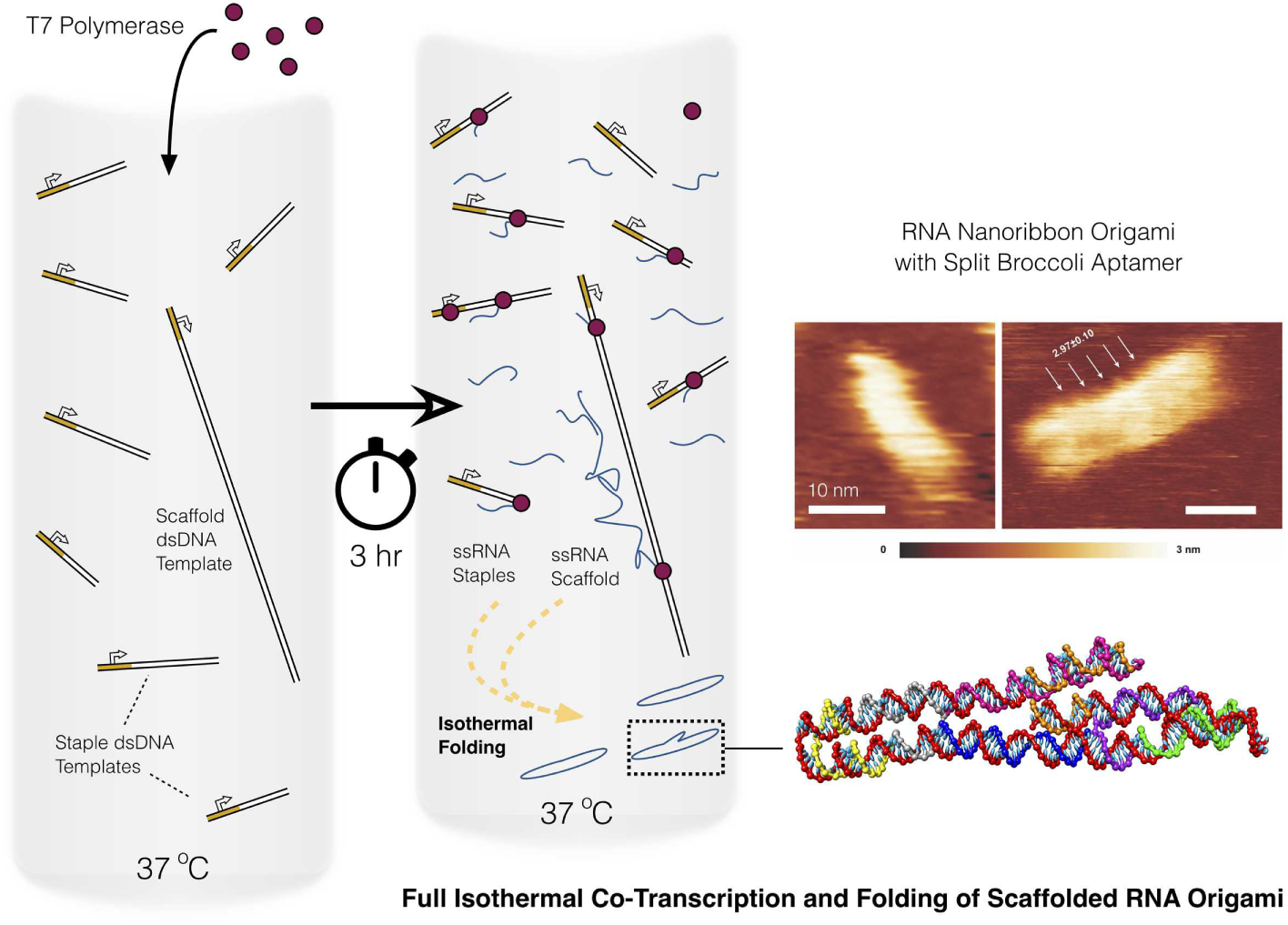

## 1 Introduction

RNA plays sophisticated roles with different and essential coding and noncoding functions. mRNAs, tRNAs, rybozymes, aptamers and CRISPR RNAs are just few examples of RNA species in a vast functional repertoire. Considering the diversity in functional and structural motifs, the emerging field of RNA nanotechnology has been developing rapidly over the past years. As a consequence, a variety of RNA nanostructures with different functionalities, sizes and shapes have been created to investigate and successfully demonstrate their potential in nanobiomedicine and synthetic biology [1-12].

Different self-assembly strategies have been adopted to design RNA nanostructures [13] ranging from RNA architectonics [14-18] to single-stranded self-assembly [19].

Furthermore, taking advantage of the sequential transcription reaction by bacteriophage RNA polymerase, single-stranded RNA origami has been synthesized from long ssRNA molecules. *In vitro* transcribed and purified RNA sequences were self-folded into hearts, rectangles and rhombus shapes adapting the ssDNA origami design strategy and considering the helical periodicity difference between DNA B-type and RNA A-type helix. Large and complex ssRNA origami were synthesized using partially complemented double-stranded RNA and parallel cross-over cohesion without limitation due to RNA kissing loop interactions [20]. However, these multikilobase ssRNA nanostructures were folded using a thermal annealing ramp gradient (from 85 °C to 25 °C), thereby limiting potential *in vivo* applications, such as the scaffolding of enzymes [21].

On the other hand, previous work showed the synthesis of smaller ssRNA origami tiles and hexagonal lattices made by annealing and/or *in vitro* co-transcriptional folding that should be compatible with *in vivo* folding when genetically encoded and expressed in cells [22]. These authors developed a strategy based on the combination of hairpins, kissing loops and “dovetail seam” to promote and stabilize the folding during the T7 RNA polymerase *in vitro* transcription reaction [22, 23]. More recently, the ssRNA 2H-AE-ST tile scaffold presented by Geary and coll. [22] was used to create an aptamer-based FRET system where RNA tile synthesis *in vivo* was demonstrated by measuring FRET outputs without a direct atomic force microscopy (AFM) visualization, due to the small construct dimension [24].

Li and coll. [25] developed a different strategy in which the design concept was similar to the approach reported above, but avoiding the use of short “dovetail seams”. The RNA nanostructures were designed based on natural motifs: the folding pathway was based on hairpin formation and tertiary interactions of unpaired residues. In detail, an RNA double-square was designed using 3-way loop observed in phi29 pRNA and 90°-kink from hepatitis C virus RNA genome. These thermodynamically stable and kinetically favourable RNA nanostructures were folded both *in vitro* and *in vivo*: nonetheless, the combination of phage and viral derived structural motifs can limit the structure design, the attachment of functional units, the creation of reconfigurable and dynamic nanostructures or their suitability for future theranostic *in vivo* applications.

In addition to architectonics, single-stranded self-assembly and single-stranded origami strategies, the scaffolded RNA origami approach is still in its infancy. Drawing from DNA origami techniques in which several short staple strands sequences promote the folding of a longer single stranded scaffold into a specific shaped structure [26], we designed and synthesised RNA origami following a similar strategy. This approach can provide several advantages such as the high synthesis yield, and the possibilities to produce reconfigurable nanostructures and to incorporate multiple and different functionalities in a precise position. Indeed, previous works demonstrated the synthesis of chemically modified and siRNAs functionalized RNA origamis following a thermal gradient annealing of RNA staple strands and scaffold [27-28].

In our previous work, we went a step further to develop a biologically afunctional (i.e. bio-orthogonal by design) RNA origami able to fold at constant temperature (37 °C) after an initial denaturation step [29], which lights-up when folding to its target configuration. Seven RNA staple strands promoted the folding of a RNA bio-orthogonal synthetic De Bruijn scaffold sequence (DBS) that does not contain genetic information, restriction enzyme sites or reduced ambiguity in the addressability [30]. We demonstrated the possibility to combine scaffold bio-orthogonality, physiologically compatible folding and assembly monitoring using a new RNA-based reporter system. The folding was monitored using our new split Broccoli aptamer system [29].

Motivated by our previous study, and with an ultimate goal of enabling RNA origami expression in living cells, here we demonstrate a full isothermal protocol for scaffolded RNA origami assembly via co-transcriptional folding. We maintain the design simplicity and the scaffold nucleotide composition, while changing each staple in order to guarantee a reasonable yield of the desired transcripts and a low aberrant products synthesis during the *in vitro* transcription by T7 RNA polymerase. The RNA origami is co-transcriptionally folded into a nanoribbon shape at 37 °C: in detail, during the scaffolded RNA assembly, scaffold and staple strands are *in vitro* transcribed and folded in a one-pot reaction. The RNA origami assembly is verified by gel assay and well characterized by AFM. The RNA nanostructures self-assembly is also successfully confirmed and selectively detected using our split Broccoli aptamer system: the transcription mix is analyzed by in-gel imaging and the tagged RNA origami shows a clear fluorescent band. Finally, the assembled RNA origami nanoribbon is visualized and simulated at equilibrium using the oxRNA coarse-grained model.

The *in vitro* transcribed and folded RNA origami described here can be compatible with expression in bacterial cells and its self-assembly can be monitored with a protein-free fluorescence detection system using an in-gel imaging assay as a rapid and specific pre-screening method.

## 2 Experimental

### 2.1 Materials and reagents

(5*Z*)-5[(3,5-Difluoro-4-hydroxyphenyl)methylene]-3,5-dihydro-2-methyl-3-(2,2,2-trifluoroethyl)-4*H*-imidazol-4-one (DFHBI-1T) was purchased from Tocris Bio-techne. A DFHBI-1T stock solution (20 mM) was prepared in DMSO, stored in the dark at – 20°C and used within 2 weeks. All RNA oligonucleotides were purchased from Eurogentec and resuspended in Ultra Pure™ distilled water to give stock solutions of 100 μM and stored at – 80 °C. All DNA oligonucleotides and gBlocks Gene Fragment were purchased from IDT. The DNA oligonucleotides were resuspended in Ultra Pure™ distilled water to give stock solutions of 100 μM and stored at – 20 °C. The gBlocks Gene Fragment was resuspended at a final concentration of 10 ng μL^−1^ and stored at – 20 °C. EDTA 0.5 M pH 8.0, Ultra Pure™ 10x TBE buffer, Ultra Pure™ 1 M Tris-HCl pH 7.5, Ultra Pure™ distilled water, 5 M NaCl (0.2 μm filtered), SYBR® Gold, dithiothreitol molecular biology grade, GlycoBlue™ Coprecipitant (15 mg/mL), NTP (100 mM each), 6% Novex™ TBE gel, 10% Novex™ TBE gel and 10% TBE-Urea gel were purchased from Thermo Fischer Scientific. Spermidine BioUltra, HEPES 1 M pH 7 Bioreagent, KCl 1 M BioUltra, MgCl_2_ 1 M BioUltra, 3-aminopropyltriethoxysilane (APTES) and agarose were purchased from Sigma-Aldrich.

### 2.2 Scaffold and staples design

The synthetic RNA scaffold and RNA staples were generated with the computer code presented by Kozyra et al. [30]. The RNA origami ribbon (Fig. S1 in the ESM) has been designed as previously described [29]: in the present study, RNA staple strands were changed (r1, l1, r2, l2, f and s1) or elongated (s2) at the 5’ end of few bases (1 up to 3 bases) in order to start with -GG or -GA (first and second nucleotides of the transcribed region) necessary for an efficient T7 transcription yield. Furthermore, all sequences were checked to reduce aberrant products synthesized during the transcription reaction. The RNA staple strands sequences and the scaffold sequence were placed downstream from the T7 promoter (5’-TTCTAATACGACTCACTATA-3’). The dsDNA templates for the transcription were two annealed oligonucleotides (for staple strands transcription) or a gBlocks Gene Fragment sequence (for scaffold transcription): all the dsDNA templates carried the T7 promoter sequence and the template to be transcribed (Table S1 in the ESM).

### 2.3 OxRNA simulation

We simulated the RNA ribbon/split aptamer system using a nucleotide-level coarse-grained RNA model, oxRNA [31, 32]. In particular, we investigated thermodynamic stability of a pre-assembled nanoribbon in isothermal conditions over a period of time.

In order to simulate the system, the caDNAno design file [33] were used in combination with caDNAno interface scripts (available with oxRNA software) to generate the initial topology of the simulated system. Preceding the simulation, the system was first relaxed, and the interaction type changed to RNA. To form initial configuration mutual traps were used according to the software documentation. Briefly, a harmonic force was introduced to pull selected particles from one strand of split aptamer toward reference particles on the other strand. This process used low stiffness parameter (0.1) until a predefined equilibrium distance of the trap was reached, in this case, 1.5 simulation units of length, corresponding roughly to the hydrogen bonding potential. A short simulation was run, and after the equilibrium was reached and the split aptamer structure formed, the mutual traps were removed.

The system was then simulated over a longer time period and the trajectory file recorded. Molecular Dynamics (MD) was the selected simulation algorithm and the simulation was run for 2*10^7 simulation steps which, in physical units, corresponds to 61.2 μs. The temperature of the system was set to 310K (∼37 °C). The thermostat used was john thermostat, which is the optimal thermostat that emulates Brownian dynamics. The john thermostat, which is an Andersen-like thermostat, was used since it is the optimal thermostat that emulates Brownian dynamics [34].

### 2.4 RNA scaffold synthesis and purification

Double-stranded gBlocks Gene Fragment containing T7 promoter (gBlocks DBS scaffold, Table S1 in the ESM) was amplified using Phusion® DNA polymerase (NEB) and DBS forward/DBS reverse primers (Table S1 in the ESM), as previously described. Briefly, an initial denaturation at 98 °C for 30 sec was followed by 15 cycles of denaturation at 98 °C for 10 sec, annealing at 60 °C for 20 sec and extension at 72 °C for 15 sec. Finally, an additional extension was achieved for 5 min at 72 °C. The PCR product was purified using Monarch® PCR & DNA Cleanup kit (NEB) and the DNA concentration was measured on a NanoDrop spectrophotometer. The size of purified amplicon was evaluated on 1.5% agarose gel in TBE for 1 h 40 min at 110 V: the gel was pre stained with Nancy-520 and visualized under UV illumination. The low molecular weight DNA ladder (NEB) was used as molecular weight marker.

The purified template was transcribed *in vitro* at 37 °C for 1 h and 30 min using Ampliscribe™ T7-Flash™ Transcription kit (Epicentre). After DNase treatment at 37 °C for 15 min, the RNA transcript was purifed using RNA Clean & Concentrator™ (Zymo Research), quantified using a NanoDrop spectrophotometer and used as scaffold sequence for the RNA origami assembly reaction.

Alternatively, the amplified and purified scaffold template was used for co-transcriptional folding.

### 2.5 Synthetic RNA origami nanoribbon folding and native PAGE

The set of RNA staple strands [29] were mixed in 10-fold excess with RNA scaffold in 50 μL of folding buffer (10 mM MgCl_2_, 20 mM Tris-HCl pH 7.5, 1 mM EDTA pH 8.0, [27]). After an initial thermal denaturation step at 75 °C for 1 min and a snap cooling (−1 °C/0.42 sec), the mixture was subjected to a folding step at 37 °C for 20 min. Folding was performed with a SensoQuest Labcycler GeneFlow thermalcycler. Samples were run on 6% Novex™ TBE gel in 1x TBE buffer at 100 V for 40 min at low temperature (below 10 °C). After staining with SYBR® Gold in 1x TBE for 5 min, the gels were visualized using Typhoon laser scanner and ImageQuant TL software (normal sensitivity; GE Healthcare Life Sciences). The low range ssRNA ladder (NEB) was used as molecular weight marker.

### 2.6 Double-stranded DNA template annealing, purification and transcription

RNA staples to be transcribed are less than 100 bases, ranging from 26 to 51 nucleotides. For this reason, DNA templates for *in vitro* transcription were obtained from two annealed single-stranded DNA oligonucleotides for each staple strand sequence: complementary sense and antisense strands (RP-HPLC or PAGE purified) containing the T7 promoter sequence and the RNA sequence to be transcribed were annealed by incubating equimolar concentrations (5 μM) of forward and reverse strand in TE supplemented with 12.5 mM MgCl_2_ (r1, r2, l1 and l2) or with 50 mM NaCl (s1, s2 and f). The samples were heated at 95 °C for 2 min and slowly cooled down at 25 °C (72 cycles, 38 sec/cycle, -1°C/cycle): all annealing processes were performed with a Biometra TRIO Analitik jena thermalcycler. Samples were loaded and run on 10% Novex™ TBE gel in 1x Tris borate EDTA (TBE) buffer at 200 V for 45 min. After staining with SYBR® Gold in 1x TBE for 5 min, the gels were visualized using Typhoon laser scanner and Image Quant TL software (normal sensitivity; GE Healthcare Life Sciences). The low molecular weight DNA ladder (NEB) was used as molecular weight marker. Annealed dsDNA were purified from polyacrylamide gels by the ‘crush and soak’ method, as previously described [35] with some modifications. The bands of interest were cut out and the gel slices were transfered to a 0.5 mL tube that was pierced with a 20-G needle and placed inside a 1.5 mL microcentrifuge tube. The tube was centrifugated at 20000g for 3 min to force the gel through the needle hole. 0.67 mL of DNA soaking buffer (0.3 M NaCl, 10 mM Tris-HCl pH 7.5, 0.97 mM EDTA) were added to the recovered gel and incubated overnight with agitation at room temperature. Gel and soaking buffer were purified from gel debris using a Freeze’n Squeeze DNA gel extraction spin column (Bio-Rad) that was centrifugated at 20000g for 3 min at 4 °C to recover the soaking mixture. The eluted DNA was transferred to a fresh microcentrifuge tube and precipitated at least 1 hour at -20 °C using 0.68 mL isopropanol supplemented with 1 μL GlycoBlue™ Coprecipitant. After centrifugation at 4 °C (30 min at 20000g), the pellet was washed in 0.75 mL of 80% ice-cold ethanol, air-dried and resuspended in 12 μL of 10 mM Tris-HCl pH 7.5. DNA samples were run on 10% Novex™ TBE gel and visualized as described above. Purified and unpurified samples were measured using NanoDrop One/One^C^ spectrophotometer (Thermo Scientific).

Each purified or unpurified dsDNA template encoding the RNA staples strands were used for RNA synthesis. Transcription reactions were performed in 20 mM Tris-HCl pH 7.6, 10 mM MgCl_2_, 2.5 mM of each rNTPs, 4 mM DTT, 2 mM spermidine, 2 units/μL T7 RNA polymerase (NEB). The transcription mix was incubated at 37 °C for 3 h and treated with 1 μL DNase RNase free (NEB) at 37 °C for 15 min. After dilution, samples were run on 10% Novex™ TBE gel in 1x TBE buffer at 200 V for 45 min. After staining with SYBR® Gold in 1x TBE for 5 min, the gels were visualized using Typhoon laser scanner and ImageQuant TL software (normal sensitivity; GE Healthcare Life Sciences). The low range ssRNA ladder (NEB) and ZR small-RNA™ ladder (Cambridge Bioscience) were used as molecular weight marker.

### 2.7 Co-transcriptional folding of RNA origami nanoribbon

The concentrations of each dsDNA template encoding the RNA staples strands and scaffold were measured using NanoDrop One/One^C^ spectrophotometer (average concentration values were calculated from 3 measurements for each sample). Different molar concentrations of DNA templates were used for RNA synthesis and RNA origami folding during the T7 transcription: 5 nM gBlock Gene Fragment (scaffold), 5 nM of each s1, s2 and f dsDNA, 10 nM of each r1 and l2 dsDNA, 20 nM l1 dsDNA, 30 nM r2 dsDNA. Transcription reaction was performed in 200 mM Tris-HCl pH 7.6, 10 mM MgCl_2_, 2.5 mM of each rNTPs, 4 mM DTT, 2 mM spermidine, 2 units/μL T7 RNA polymerase (NEB). The transcription mix was incubated at 37 °C for 3 h and treated with 1 μL DNase RNase free (NEB) at 37 °C for 15 min.

### 2.8 Co-transcribed RNA origami: native PAGE and in-gel imaging

After dilution, samples were run on 6% Novex™ TBE gel in 1x TBE buffer at 100 V for 40 min at low temperature (below 10 °C). After staining with SYBR® Gold in 1x TBE for 5 min, the gels were visualized using Typhoon laser scanner and ImageQuant TL software (normal sensitivity; GE Healthcare Life Sciences). The low range ssRNA ladder (NEB) was used as molecular weight marker.

To confirm the RNA origami self-assembly by incorporation of Split Broccoli aptamer system, in-gel imaging with fluorophore DFHBI-1T [36] was performed with some modifications. Briefly, RNA origami sample, partially folded samples and Broccoli aptamer (prepared as previously described, [29]) were loaded in the polyacrylamide gels. The gels were washed three times for 5 min in RNase free water and then stained for 20-25 min in aptamer buffer containing 1.26 μM DFHBI-1T, 40 mM HEPES pH 7.4, 100 mM KCl, 1 mM MgCl_2_. The gels were imaged using Typhoon laser scanner (excitation 488 nm, emission 532 nm): bands were analyzed using ImageQuant TL software. Then, the gels were washed three times with Ultra Pure™ distilled water, stained with SYBR® Gold in 1x TBE for 8 min, and visualized using Typhoon laser scanner. The low range ssRNA ladder (NEB) was used as molecular weight marker.

### 2.9 Atomic Force Microscope (AFM) imaging

All the experiments were conducted on a commercial AFM Cypher ES (Asylum Research, Oxford Instruments, Santa Barbara, CA) and with the tip and cantilever fully immersed into the liquid. The vertical oscillation of the tip was controlled by photothermal excitation (Blue Drive) and the experiments were conducted at 25.0 ± 0.1 °C. All measurements, were conducted in Amplitude Modulation (AM-AFM) mode using SCANASYST-Fluid+ (Bruker, Camarillo, CA) cantilever and with a setpoint ratio between the free amplitude and imaging amplitude of ∼0.8.

Freshly cleaved mica was passivated for 5 min with 10 μL of 0.1% APTES in water to ensure the adhesion of the negative charged RNA origami structures on the negative mica surface. After three washing steps with 10 mM MgCl_2_, 20 mM Tris-HCl pH 7.5, 1 mM EDTA pH 8.0, the transcription reactions were diluted 1:100 in the same buffer, added (2 μL) to the passivated mica surface, allowed to adsorb for 5 min in a chamber and imaged immediately. When purified RNA origami samples were imaged, 8 μL of sample were added to the passivated mica surface and allowed to adsorb for 5 min in a chamber as above.

All the images were corrected for tilt (line or plane fattening) and lightly low-pass filtered to remove grainy noise using the WSxM sofware (Nanotec Electronica, Madrid, Spain) [37].

## 3 Results and Discussions

### 3.1 Staple strands and RNA origami design

Recently, we designed a 212 nt biologically inert (i.e. bio-orthogonal) and uniquely addressable De Bruijn scaffold sequence (DBS) characterized by lack of genetic information, restriction enzyme sites and reduced ambiguity in the addressability: the bio-orthogonality of the RNA scaffold designed *in silico* was demonstrated in *E. coli* cells [29].

Here, we used our previous optimized RNA scaffold and a new set of seven staple strands starting with -GG or -GA nucleotides at the 5’ end (Fig. S1 and Table S1 in the ESM) in order to promote a reasonable transcriptional yield and allow an efficient control of the 5’ sequence content [38]. In detail, transcription reactions of each RNA sequence were carried out with T7 RNA polymerase and synthetic DNA containing the T7 promoter: the above mentioned 5’ end starting nucleotides can guarantee efficient transcription and efficient control of the 5’ sequence content during the T7 polymerase reaction [38].

Furthermore, although T7 RNA polymerase is a highly specific enzyme, the desired short RNA is usually accompanied by undesired products that are longer or shorter than the expected transcript [39-44]. Besides its higher DNA affinity, T7 RNA polymerase is able to synthesize RNA from single or double-stranded RNA template [42, 43]. It has been demonstrated that RNA extension can occur if 3’ end has a self-complementarity (*trans* mechanism) or folds back on itself in *cis* to form extendible duplexes [40, 42, 43]. Next generation RNA-Seq analysis has demonstrated that primer extension occurs predominantly via a *cis* self-primer mechanism and strongly depends on pairing A at 3’ end with the U, 9 bases upstream [43]: as a result, partially double-stranded RNA byproducts are synthesized. In order to reduce the amount of longer undesired RNA extension products catalyzed by T7 RNA polymerase, all staple strands sequences were examined to take account of the 3’ end self-complementarity and pairing between 3’ end adenine and uracil, 9 bases upstream: the analysis revealed that all RNA sequences were lacking of these distinct 3’ end characteristics. In addition, false transcriptions are mainly observed when the correct product is free of stable secondary structures at the 3’ end: the aberrant transcripts are longer than the coded RNA [40]. It has been reported that improved transcription of the correct length RNA can be obtained by adding hairpins to the 3’ and 5’ end: these hairpins decrease the transcription of incorrect-lenght products, but still a series of bands can remain [45, 46]. In our work, staple sequences were optimized to weaken secondary structures and avoid hairpin formation [30].

### 3.2 DNA template preparation and transcription

The scaffold sequence and each of the staple strands sequences were placed down-stream from the T7 RNA polymerase promoter and the linear double-stranded DNA templates were characterized by blunt ends in order to reduce the synthesis of spurious transcripts.

Scaffold DNA template was purified from an amplified gBlocks Gene Fragment (Fig. S2 in the ESM). As staple strands to be transcribed were shorter (ranging from 26 to 51 bases), DNA templates were obtained from an annealing step of two RP-HPLC or PAGE purified oligonucleotides.

Annealed DNA templates corresponding to staple strands showed multiple bands (Fig. S3 in the ESM) presumably due also to the presence of byproducts in addition to the desired chemically synthesized oligonucleotide (RP-HPLC and PAGE purification usually can provide a purity level up to 85% and around 85-90%, respectively).

To increase the purity, annealed DNA templates were purified from polyacrylamide gel by the ‘crush and soak’ method using GlycoBlue™ as coprecipitant to facilitate the DNA recovery and reach a concentration compatible with the subsequent transcription reaction avoiding any mixture dilution. All samples were run and checked on native PAGE (Fig. S4 in the ESM).

The use of GlycoBlue™ coprecipitant at a final concentrations (22 μg/mL) below the recommended value (50-150 μg/mL) did not allow an accurate DNA quantification due to the blue dye absorbance and the low A260/A230 ratio value (0.8). Furthermore, there were no differences in the transcription products when purified and unpurified dsDNA samples were used as template. For these reasons, we decided to *in vitro* transcribe unpurified DNA templates.

RNA sequences were transcribed *in vitro* and analyzed by denaturing PAGE. In order to improve the transcription, different concentrations of DTT and spermidine were tested: transcription of each staple strand DNA template in 4 mM DTT and 2 mM spermidine showed a distinct main band corresponding to the full-lenght transcript (Fig. S5 in the ESM). In addition to the full-length transcript, the synthesis of a pattern of products with a higher molecular weight was also observed as previously described [40, 46]. Triana-Alonso and coll. [40] demonstrated the synthesis of longer transcript as a secondary process following an induction time which is different for every template: a minimal concentration of transcript is required for aberrant transcripts that accumulated at the end of long incubation as a result of a late event during the transcription. As RNA staple strands were designed to not support RNA-primed RNA extension, we hypothesized that, after the production of a certain amount of transcript, T7 polymerase accepted template RNA sequences that were not folded in 3’ end stable secondary structures. We then assumed that during the co-transcriptional folding, duplex formation related to the RNA origami folding can sequester the staple strands 3’ end reducing RNA-templated extension and undesired byproducts.

Finally, abortive synthesis was observed in T7 RNA polymerase transcription as a fundamental and major feature of early stages: several short transcripts were produced (maximum lenght of 12 nt) due to a kinetic competition between dissociation of the enzyme-DNA-RNA complex and incorporation of successive nucleotides [47]. During the staple strands transcription, low amount of abortive products (Fig. S4 in the ESM) were produced without influencing the transcription efficiency. Previous studies showed that both abortive products and intermediate products were observed as an unavoidable result of the *in vitro* transcription [41, 47].

### 3.3 OxRNA simulation

The simulated RNA ribbon/split aptamer system stayed at equilibrium and the split aptamer structure was preserved at 37 °C. The two bulge loops (3 nt and 2nt-long, respectively) were visible in the aptamer structure (Fig. 1, red arrows). The hybridisation between split aptamer strands and the origami scaffold allowed for a degree of flexibility between the aptamer and the ribbon. Interestingly, the simulation suggested that the internal loop (located between the red arrows) formed a double-helical structure despite the mismatched nucleotides. However, the Gibbs’s free energy of that interaction was higher than a typical complementary sequence of similar length and thus less stable.

**Figure 1.**
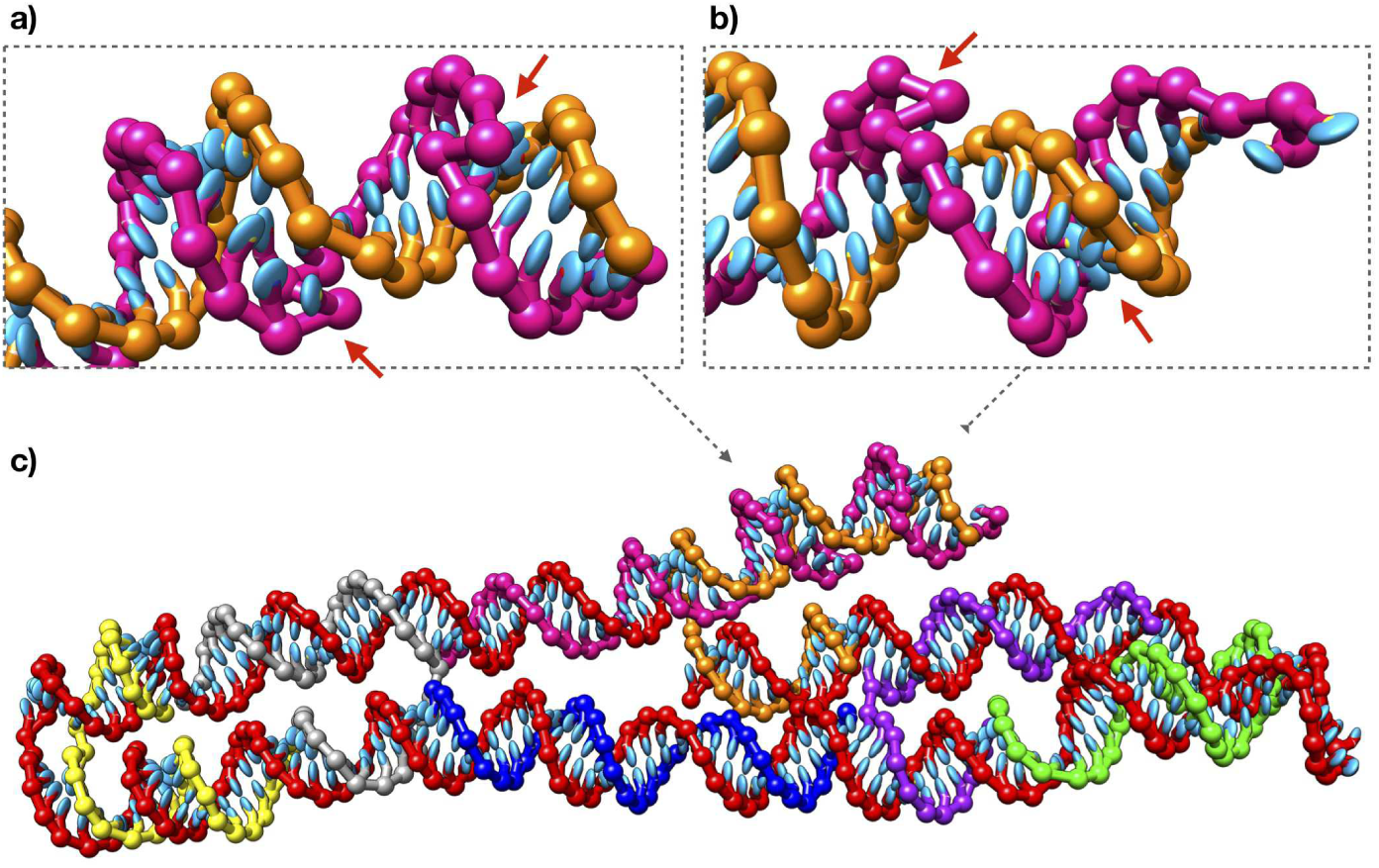
oxRNA simulation of a pre-assembled RNA ribbon. Final configuration of the simulated system is shown after an equilibration period. The RNA ribbon is formed of the scaffold (red) which is bound by 5 staples (various colours). The two split aptamer strands (pink and orange) are located at the 5’ and 3’ ends of the scaffold, accordingly. Enlarged views show the front (a) and the back (b) of the aptamer structure in respect to the RNA ribbon (c). Red arrows indicate the location of bulge loops.

### 3.4 RNA origami co-transcriptional folding and AFM imaging

Self-assembly at physiologically compatible temperature can expand and improve the RNA nanotechnology applications under intracellular conditions. To date, the ability to design RNA nanostructures that co-transcriptional fold into the target shape has been demonstrated using a single-stranded origami approach [22] or a multiple sequences self-assembly technique [19, 48, 49]. In detail, RNA tile nanostructures can fold from a single-stranded RNA sequence [22], while RNA nanocubes can assemble from optimized short strand sequences [19, 49].

In contrast to the above strategies, co-transcriptional folding of a scaffold strand using complementary staple strands has not yet been demonstrated. Previous works showed RNA origami self-assembly by thermal annealing in a linear temperature ramp, thereby limiting potential *in vivo* applications [27, 28].

In our recent work, we reported the successful bio-orthogonal RNA origami isothermal folding at 37 °C with a prior denaturation step at 75 °C [29]. As a proof of principle, here we demonstrate the co-transcriptional folding of a genetically encoded scaffolded RNA origami using our previously designed bio-orthogonal DBS scaffold. Double-stranded DNA templates encoding scaffold and staple strands were transcribed by T7 RNA polymerase in an optimized transcription buffer containing 4 mM DTT and 2 mM spermidine. Furthermore, partially folded scaffold samples were transcribed using two subsets of staples consisting of Staple s1 and s2, and Staple s1, s2, l1 and r1. These subsets were considered not only to demonstrate their different electrophoretic migration pattern, but also their different fluorescence band intensity in the following in-gel imaging experiments.

After a DNase treatment, the unpurified transcription mixtures were directly analyzed by nondenaturing polyacrylamide gel electrophoresis: gel image showed a distinct band around the expected size (black arrow in Fig. 2) with a different migration distance compared to that of partially folded products, suggesting the correct folding. Bands corresponding to scaffold and partially folded products were almost not present in the RNA origami sample indicating a high folding efficiency.

**Figure 2.**
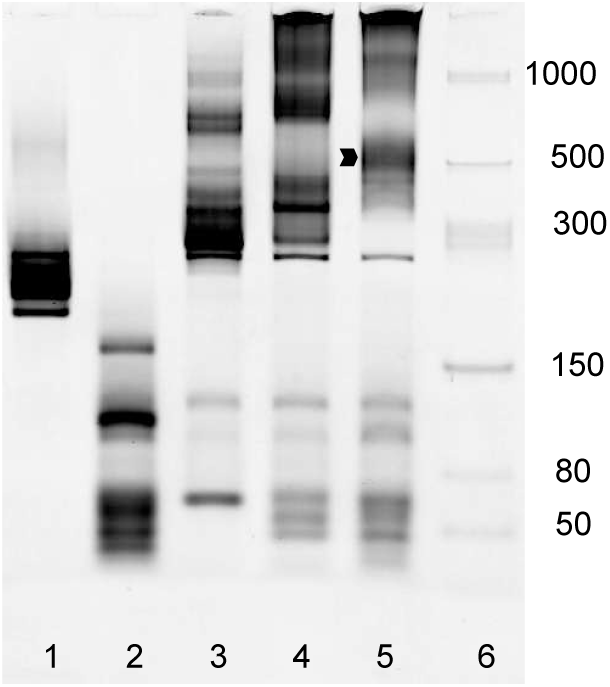
6% TBE gel electrophoresis of transcription products after SYBR® Gold staining. Lanes. 1: transcribed RNA scaffold; 2: transcribed RNA staple strands; 3: transcribed RNA scaffold, Staples s1 and s2; 4: transcribed RNA scaffold and RNA Staples s1, s2, r1 and l1; 5: co-transcriptional folded RNA (black arrow); 6: low range ssRNA ladder. Molecular size in nucleotides are indicated.

The co-transcriptional folded nanostructure band showed a slight difference in the migration distance when compared to nanostructures assembled from synthetic RNA staple sequences (Fig. S6 in the ESM), as previously observed [19]: the electrophoretic migration of unpurified transcribed samples was influenced by the different and more complex reaction mixture composition [50]. Finally, we found a higher band closed to the well (Fig. 2): as a result of both thermal ramp or co-transcriptional folding, higher bands can appear [19, 22, 28, 51] and can correspond to aggregates [28], spurious assemblies [51] or undesired kinetically trapped products [22].

In detail, abortive and elongated RNA products of incorrect length are an unavoidable result of the *in vitro* transcription [41, 47] and they can contribute to the formation of spurious assemblies [51]. Stewart and coll. [51] showed that despite the presence of these transcription by-products able to generate unknown folding, the one-pot method yields the desired assemblies. The latter authors showed example of gel electrophoresis images with significant amount of abortive and elongated products. Furthermore, AFM images of tubular assemblies and flat lattices revealed a ‘noisy’ environment. In the perspective of future *in vivo* isothermal folding, it should be noted that aberrant products produced *in vitro* by T7 RNA polymerase are not commonly synthesised inside living cells [40].

The unpurified RNA origami sample was characterized by AFM immediately after the co-transcriptional folding. Samples were diluted and deposited on a passivated mica surface using a silane solution instead of Mg^2+^. Indeed, it has been suggested that the different phosphate groups orientation in dsRNA compared to dsDNA can be a possible reason for the difficult dsRNA adsorption on mica using magnesium ions [52]. The estimated RNA nanostructures dimensions were approx. 27 nm x 5 nm, as the A-form RNA helix revealed a rise per base pair of 0.28 nm [52]. AFM images confirmed the correct folding of the co-transcribed RNA nanostructures with average lengths of 27.2 ± 3.5 nm and 8.4 ± 1.8 nm (Fig. 3 and Fig. S7 in the ESM): these images were compatible to that of purified sample (Fig. 4) previously well characterized [29], indicating that the RNA scaffold and staple strands folded into the desired shape also during the transcription reaction. RNA nanostructures showed a double-stranded RNA mean periodicity of 3.1 ± 0.3 nm (Fig. 3, right): this measurement was consistent with the A-form helical pitch of dsRNA previously reported [52]. The split Broccoli aptamer was sometimes visible as a slightly elevated side on the origami surface (Fig. 3, right) or as a thin protrusion (Fig. S8 in the ESM) depending on the orientation during the deposition on the mica surface. It has been already noted that not all the modifications of the hexagonal mini-lattices with aptamers are well visible by AFM imaging due to the extension orientation with respect to the scanning direction [53].

**Figure 3.**
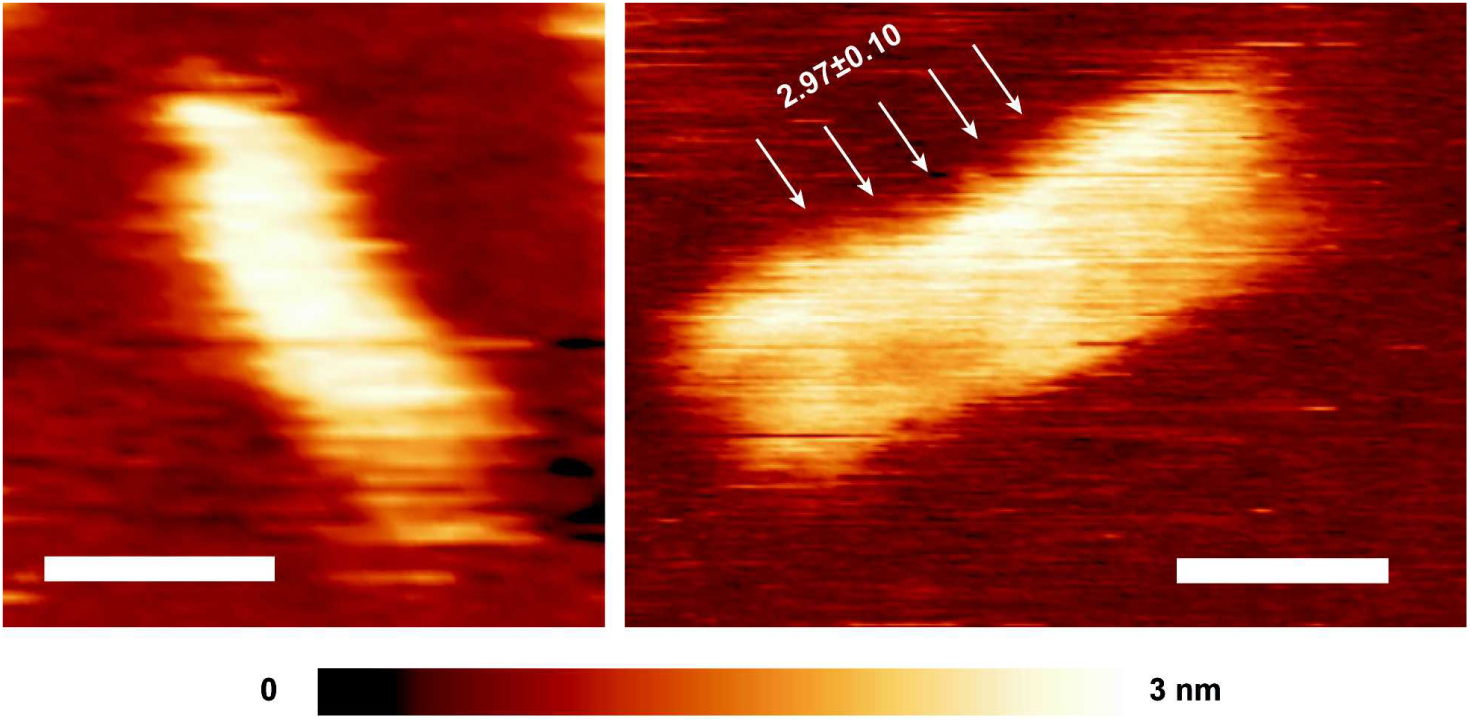
High-resolution AFM images of co-transcriptional folded RNA origami. The images were taken in solution immediately after co-transcription. The white arrows show the dsRNA helical pitch (3.2 ± 0.3 nm, [52]). Scale bar: 10 nm.

**Figure 4.**
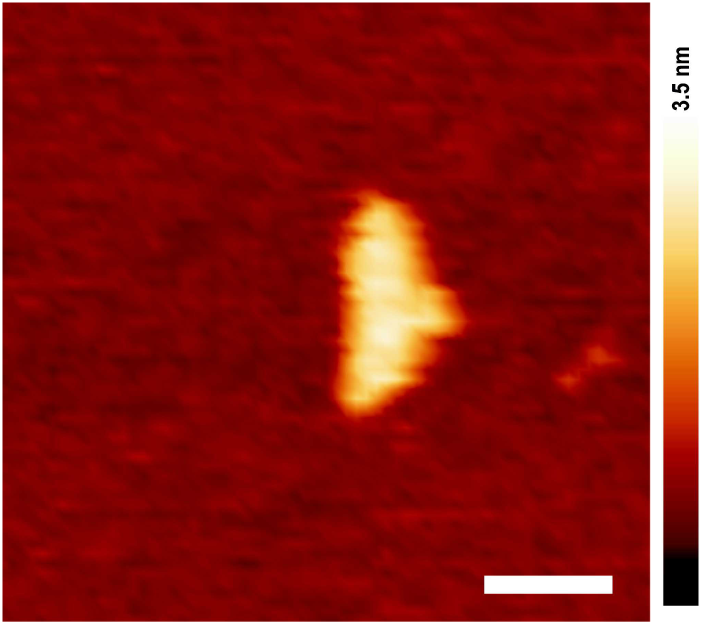
High-resolution AFM image of synthetic folded purified RNA origami. Scale bar: 20 nm.

RNA nanostructure interactions were also imaged by AFM (Fig. S8 in the ESM), presumably due to interaction between left (5’-UGUUAUACAG-3’) and right (5’-CCGGUCCGGC-3’) ssRNA scaffold regions, as previously noted [29]. Finally, smaller nanostructures were imaged as the result of a fragmentation during sample deposition and a disruption by AFM probes [25, 54].

### 3.5 Light-up RNA origami and in-gel imaging

Light-up RNA aptamers have been selected to induce a fluorescence emission upon binding specific small molecules allowing a protein-free RNA tagging [55]. Since their development, fluorescent aptamers have been used *in vitro* and *in vivo* as a valid alternative to other imaging strategies (e.g. fluorescent *in situ* hybridization and molecular beacons, [55, 56]). Malachite green, Mango, Spinach and Broccoli aptamers have been inserted in the design of different RNA nanostructures to successfully demonstrate the self-assembly and folding through their functional activation and fluorescence emission [19, 24, 29, 53]. Considering characteristics such as the short size, the robust folding under physiological conditions and the specificity for a noncytotoxic and cell permeable dye, Broccoli aptamer [57] was recently chosen to functionalize an RNA origami nanoribbon [29]. In detail, we described a new split Broccoli aptamer system that enabled us to monitor the RNA origami folding. The Broccoli aptamer was divided into two nonfunctional halves each of which was elongated in 5’ or 3’ end with two sequences complementary to the bio-orthogonal RNA scaffold. When the RNA origami self-assembled, the split sequences, called Split s1 and Split s2, were in closed proximity turning on the fluorescence [29].

Here, Staple s1 and Staple s2 were included in the RNA origami design, modified in order to start respectively with -GA and -GG, and placed downstream to the T7 promoter, as described above. After co-transcriptional folding, unpurified partially assembled scaffold and RNA origami samples were diluted, resolved by PAGE and analyzed by in-gel imaging in order to monitor the self-assembly well-characterized by AFM imaging. In detail, after PAGE and three washing steps, the gel was stained with DFHBI-1T which selectively binds Broccoli aptamer [36]. The DFHBI-1T concentration (1.26 μM) and the staining time (20-25 minutes) were optimized in order to reduce undesired fluorescent background related to the loading of unpurified transcribed samples instead of samples purified and concentrated through ultrafiltration [29]. Fluorescent imaging of the gel showed a prominent bright band with the largest peak area (approx. 6 times higher than each of the two lower bands, lane 7 in Fig. 5) corresponding to the well folded RNA nanostructures band revealed by UV imaging after SYBR® Gold staining. Lanes 5 and 6 (Fig. 5) showed partially folded scaffold with low fluorescent bands which were not present in lane 7 corresponding to the full assembled origami. In the case of synthetic partially folded scaffold samples, there were no fluorescence emissions [29]: we concluded that the fluorescent bands in the *in vitro* transcription case corresponded to unknown transcribed sequences hybridized to the scaffold.

**Figure 5.**
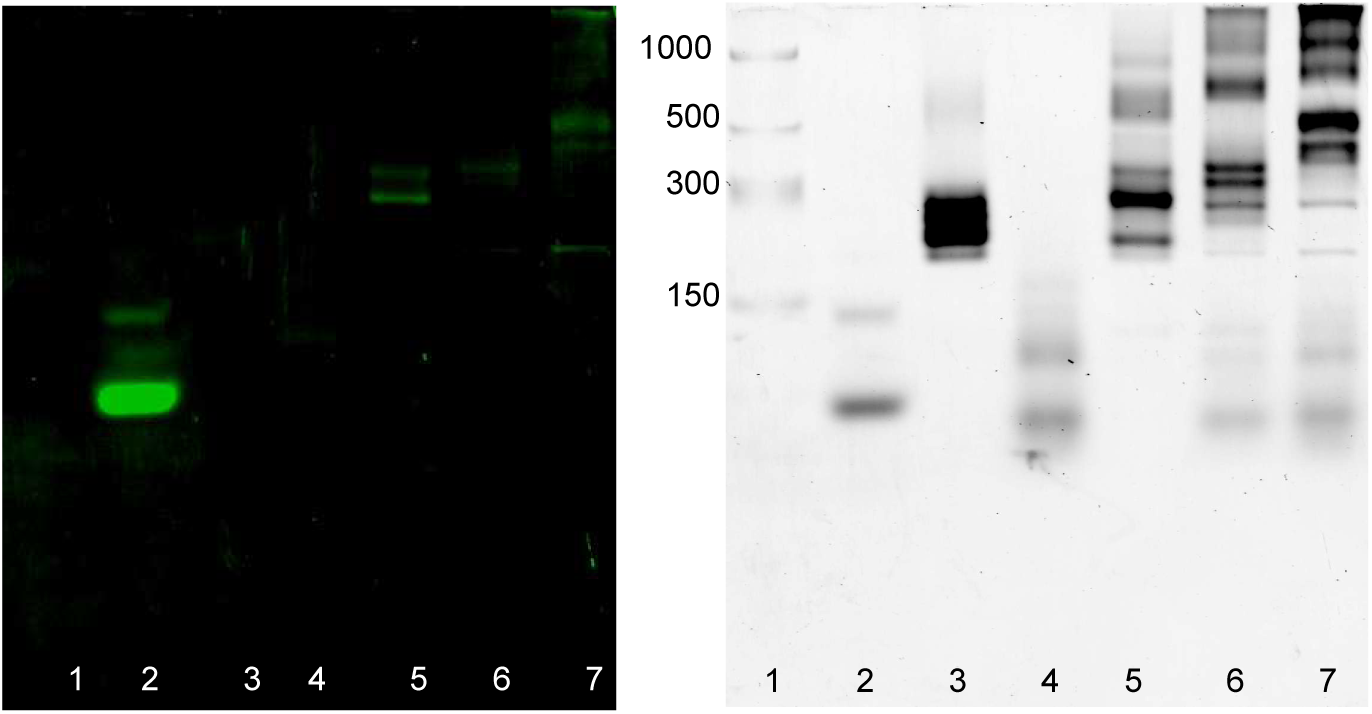
In-gel imaging of co-transcriptional folded light-up RNA origami. 10% TBE gel electrophoresis after DFHBI-1T (a) and after SYBR® Gold (b) staining. The gel was stained with DFHBI-1T for 25 min to visualize Broccoli aptamer (positive control) and co-transcriptional folded RNA origami. After 3 washing steps, the gel was stained with SYBR® Gold for 5 min to detect transcribed RNA. Lanes: 1: low range ssRNA ladder; 2: Broccoli aptamer; 3: transcribed RNA scaffold; 4: transcribed RNA staples; 5: transcribed scaffold, s1 and s2 staples; 6: transcribed scaffold, s1, s2, l1 and r1 staples; 7: transcribed RNA origami. Molecular size in nucleotides are indicated.

The double staining allowed us to confirm the correct assembly considering both the specific migration distance and the fluorescence of the RNA nanostructures band.

Considering our result and the selective rapid in-gel imaging of Broccoli tagged RNA expressed in *E. coli* [36], we concluded that this simple analysis system can be used as a pre-screening method to monitor and check if genetically encoded nanostructures are expressed in bacterial cells.

## 4 Conclusions

We have demonstrated the co-transcriptional folding of a bio-orthogonal scaffolded RNA origami in a one-pot reaction, revealing that the self-assembly of scaffold and staple strands can occur also in the transcription reaction mixture at 37 °C from double-stranded templates. In alternative to other strategies previously reported (i.e. single-stranded origami [22] and strands self-assembly [19]), the scaffolded RNA origami technique can be successfully used to design and synthesize a desired nanostructure at constant, physiologically compatible temperature, overcoming the use of synthetic purified RNA sequences not possible in living cells.

Split Broccoli aptamer functionalization was introduced into the RNA origami design allowing a simple and specific screening of the well folded nanostructures by in-gel imaging using a cell permeable and compatible dye. This approach can further suggest and confirm the use of our split light-up aptamer [29] as a reporter system to monitor the folding, avoiding the use of fluorescent proteins. Furthermore, the fluorogenic aptamer can be replaced with other functional sequences required for different purposes.

In conclusion, our results represent a further step toward *in vivo* self-assembly of nucleic acid nanostructures, and expand the design strategies used to synthesize nanostructures tailored to specific applications. In a wider picture, genetically encoded scaffolded RNA origami could represent a pathway to construct a “bio-circuit board” able to direct specific cell metabolic pathways through an orthogonal spatial and post transcriptional control.

## Acknowledgements

This work was supported by Engineering and Physical Sciences Research Council grant EP/N031962/1. Prof. Krasnogor is supported by a Royal Academy of Engineering Chair in Emerging Technology award. KV and LP acknowledge funding from the Biotechnology and Biological Sciences Research Council (grant BB/M024830/1).

## Electronic Supplementary Materials

Supplementary material (details on the sequences, design of the co-transcriptional folded RNA origami nanoribbon, gel images and AFM images) is available in the online version of this article at

