## Supplementary Materials for "Co-transcriptional folding of a bio-orthogonal fluorescent scaffolded RNA origami"

**Table S1** DNA and RNA sequences used in this study. The capital letters in bold or underlined in green correspond to T7 promoter and split Broccoli aptamer sequences, respectively.

| **Oligo name** | **Sequence [5'-3']** | **Length [bases]** |
|---|---|---|
| Broccoli ssDNA | TTCTAATACGACTCACTATAGGTATGTGGGAGACGGTCGGGTCCAGATA<br>TTCGTATCTGTCGAGTAGAGTGTGGGCTCCCACATAC | 86 |
| Broccoli formard primer | TTCTAATACGACTCACTATAGGTATGTGGGAG | 32 |
| Broccoli reverse primer | GTATGTGGGAGCCCACACTCTAC | 23 |
| gBlocks DBS scaffold | **TTCTAATACGACTCACTATA**GGTTCCTTCTTTTGGATCACAGTGGAGCA<br>ATATCTCGGGTGGCTAGGTTGTTATACAGCCGCGATATTTATCGGAAAG<br>TCGAGTGCAAGTACCCGGGAGACCGTAACTGAAACCCATAGAACCTTA<br>AACACGATTCGCTCACGGGGGTCACTTACCGGTCCGGCTCGTGTGCTCT<br>CAGTAAAGATGATCCGCCGATGGGGAAGTTTCGTCCA | 232 |
| DBS forward | TTCTAATACGACTCACTATAGGTTCC | 26 |
| DBS reverse | TGGACGAAACTTCCCC | 16 |
| Staple r1 | **TTCTAATACGACTCACTATA**GAGATATTGCTCCACTGTTTGCACTC | 46 |
| Staple r2 | **TTCTAATACGACTCACTATA**GACTTTCCGATAAATATCGCGGACCTAG<br>CCACCC | 54 |
| Staple s1 | **TTCTAATACGACTCACTATA**GATCCAAAAGAAGGAACCGTATGTGGGA<br>GACGGTCGGGTCCAGATA | 66 |
| Staple s2 | **TTCTAATACGACTCACTATA**GGTATCTGTCGAGTAGAGTG<br>TGGGCTCCCACATACTGGACGAAACTTCCCC | 71 |
| Staple l1 | **TTCTAATACGACTCACTATA**GAATCGTGTTTAATCGGCGGATCATCTTT<br>ACT | 52 |
| Staple l2 | **TTCTAATACGACTCACTATA**GAGAGCACACGATAAGTGACCCCCGTGA<br>GC | 50 |
| Staple f | **TTCTAATACGACTCACTATA**GGTTCTATGGGTTTCAGTTACGGTCTCCC<br>GGGTAC | 55 |

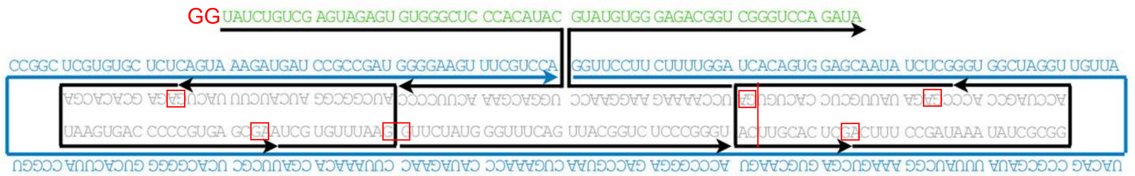


**Figure S1** The detailed design of the co-transcriptional folded RNA origami nanoribbon. The arrowheads indicate 3’ ends. Compared to the previous design [29], the staples sequences were changed in order to start with GA or GG at the 5’ end (red boxes). The RNA scaffold (in blue) was folded with seven staples strands (in black). Two staples include split Broccoli aptamer sequences (in green): two G (in red) was added to the 5’ end of staple s2.

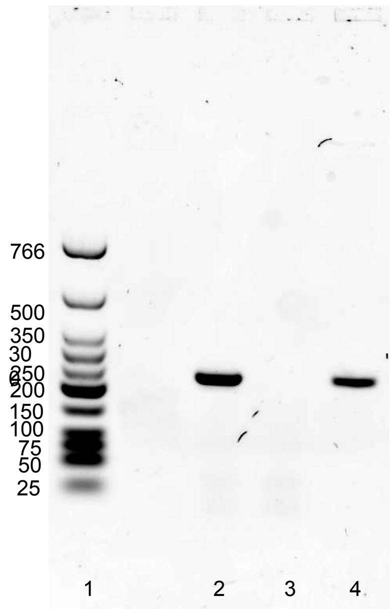


**Figure S2** 1.5% TBE agarose gel electrophoresis of amplified and purified gBlock after Nancy-520 staining. Lanes: 1: low molecular weight DNA ladder; 2: amplified gBlock; 3: blank; 4: amplified and purified gBlock using Monarch® PCR & DNA purification kit. Molecular sizes in base pairs are indicated.

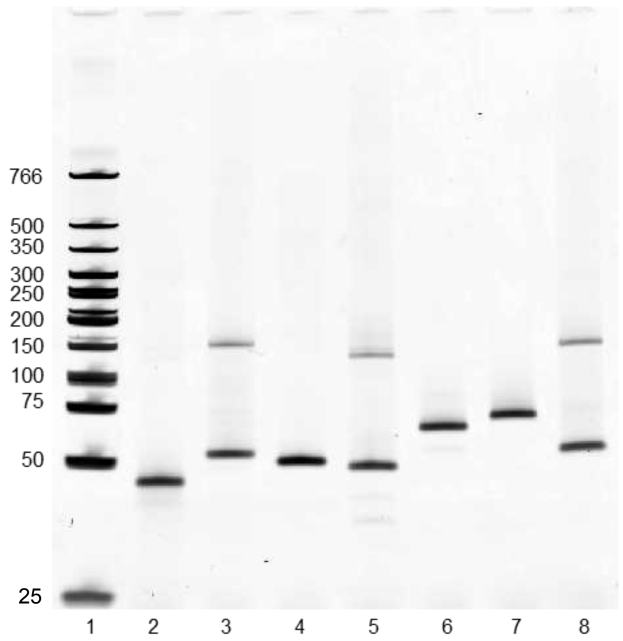


**Figure S3** 10% TBE gel electrophoresis of dsDNA samples after SYBR® Gold staining. 25 ng of dsDNA samples were loaded for each sample. Lanes: 1: low molecular weight DNA ladder; 2: dsDNA r1; 3: dsDNA r2; 4: dsDNA l1; 5: dsDNA l2; 6: dsDNA s1; 7: dsDNA s2; 8: dsDNA f. Molecular sizes in base pairs are indicated.

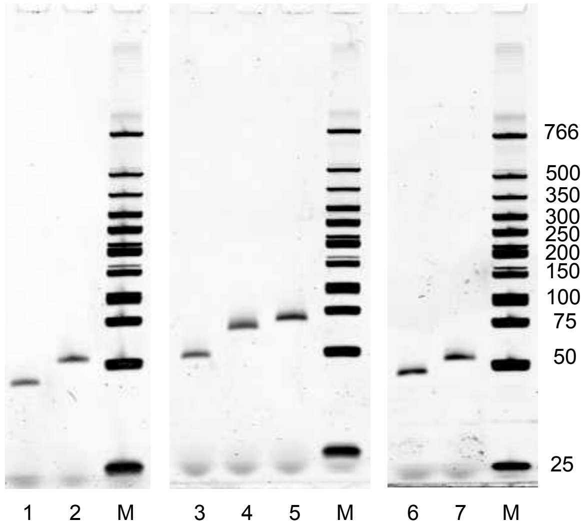


**Figure S4** 10% TBE gel electrophoresis of purified dsDNA samples after SYBR® Gold staining. All samples were PAGE purified. Lanes: M: low molecular weight DNA ladder; 1: dsDNA r1; 2: dsDNA r2; 3: dsDNA l1; 4: dsDNA s1; 5: dsDNA s2; 6: dsDNA l2; 7: dsDNA f. Molecular sizes in base pairs are indicated.

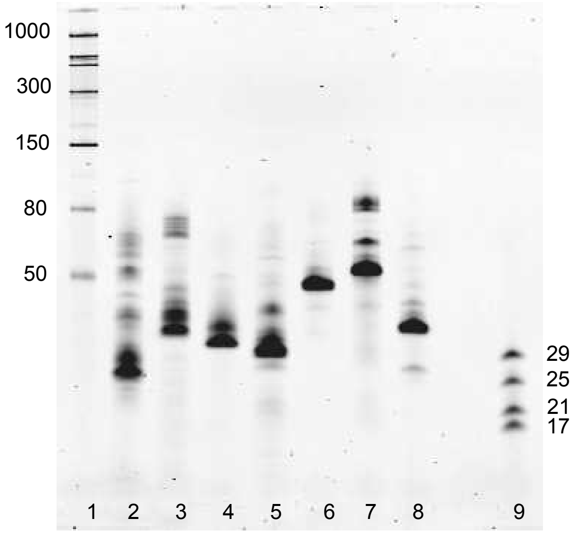


**Figure S5** 10% TBE-Urea gel electrophoresis of RNA transcripts after SYBR® Gold staining. RNA transcripts were obtained from 25 ng of dsDNA template transcribed in the transcription mixture (200 mM Tris, 10 mM $MgCl_2$, 2.5 mM NTP, 4 mM DTT, 2 mM spermidine, 100 units T7 polymerase). Lanes. 1: low range ssRNA ladder; 2: RNA staple r1; 3: RNA staple r2; 4: RNA staple l1; 5: RNA staple l2; 6: RNA staple s1; 7: RNA staple s2; 8: RNA staple f; 9: ZR small-RNATM ladder. Molecular size in nucleotides are indicated.

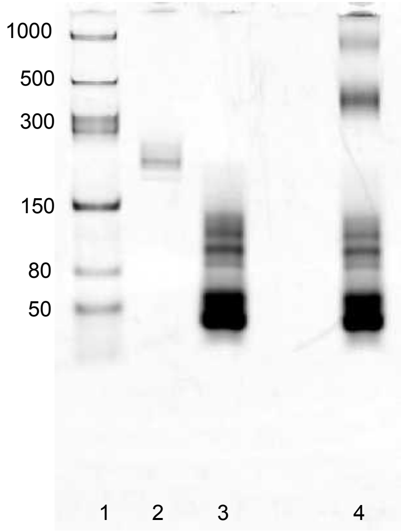


**Figure S6** 6% TBE gel electrophoresis of RNA origami sample after SYBR® Gold staining. Lanes: 1: low range ssRNA ladder; 2: RNA scaffold; 3: RNA staples; 4: RNA origami. Molecular size in nucleotides are indicated.

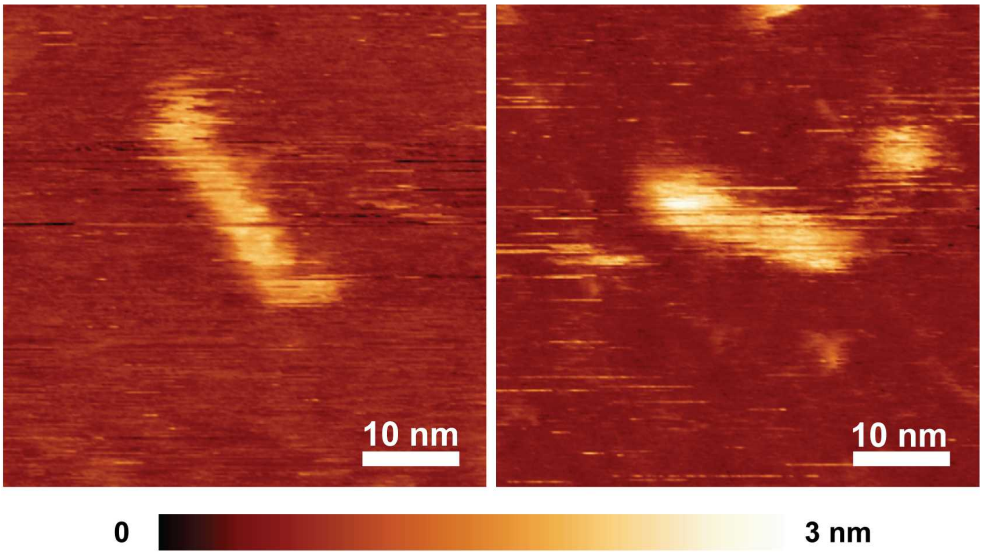


**Figure S7** High-resolution AFM images of co-transcriptional folded RNA origami. The transcription mixture was imaged immediately after preparation.

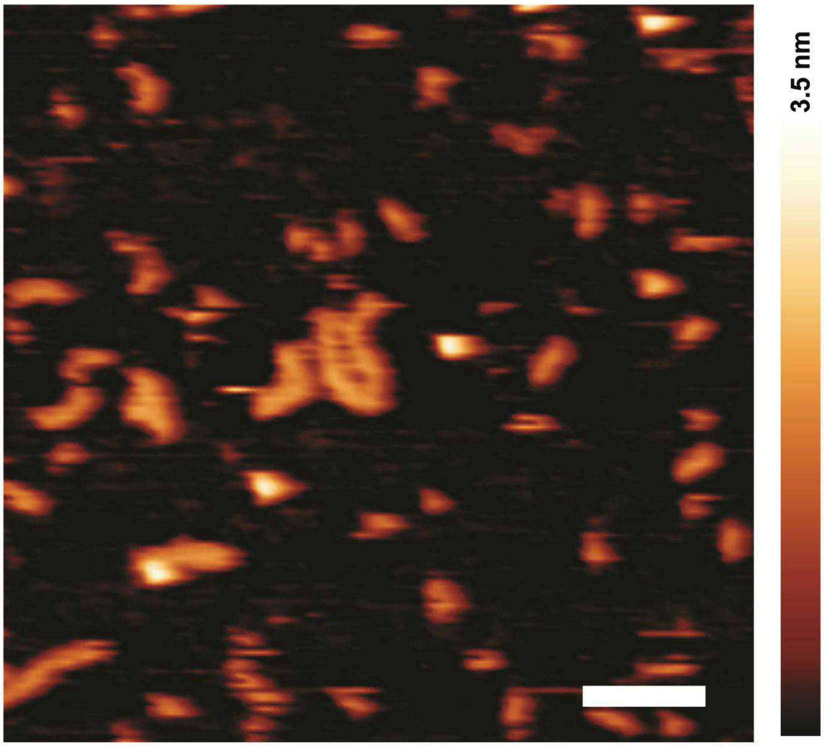


**Figure S8** AFM images of co-transcriptional folded RNA origami taken immediately after preparation and deposition on mica. Scale bar: 50 nm.
